# mTOR inhibition augments antitumor immune effector response by reprogramming the *TP53*-mutant, immune-cold HNSCC tumor microenvironment

**DOI:** 10.64898/2026.06.04.730026

**Authors:** Priyatosh Nath, Alok Khandelwal, Chun Li, Tara Moore-Medlin, Srivatsa Surya Vasudevan, Victor Albornoz Alvarez, Omar E. Franco, J. Silvio Gutkind, Cherie-Ann O. Nathan

## Abstract

**Introduction:** Resistance to immunotherapy remains a major clinical challenge in *TP53*-mutant head and neck squamous cell carcinoma (HNSCC), a disease subset characterized by immune exclusion, high recurrence, and poor outcomes. Given the constitutive activation of PI3K/AKT/mTOR signaling in *TP53*-mutant HNSCC and its role in disease progression, we investigated whether the mTOR inhibition (mTORi) could overcome immune resistance and improve outcomes.

**Methods:** We evaluated the effects of mTOR inhibitor everolimus in *TP53*-mutant, anti-PD-1-resistant syngeneic HNSCC cell lines and syngeneic tumor models. Tumor microenvironment (TME) changes, including immune cell infiltration, immune checkpoint expression, and key pathways associated with immune suppression and angiogenesis were assessed to define the mechanisms underlying TME remodeling.

**Results:** Everolimus significantly suppressed tumor growth in syngeneic HNSCC models. At the cellular level, everolimus significantly increased intratumoral CD8+ T cell and dendritic cell (DC) infiltration while reducing the accumulation of regulatory T cells (Tregs). Mechanistically, everolimus induced a cytokine/chemokine response, marked by increased TNF-α/CXCL10 expression, leading to enhanced immune infiltration. Everolimus also inhibited the HIF-1α/VEGFA angiogenic axis, a central driver of immune exclusion and myeloid-derived suppressor cell (MDSC) recruitment. Furthermore, everolimus treatment attenuated PD-1/PD-L1 signaling by reducing PD-1 and PD-L1 expression, respectively, in T cells and tumor cells, thereby restoring T-cell cytotoxic competence.

**Discussion:** These findings demonstrate that mTORi with everolimus reverses multiple mechanisms of immune resistance in *TP53*-mutant HNSCC by promoting immune cell recruitment, suppressing immunosuppressive pathways, and enhancing anti-tumor T cell activity. Collectively, these results support mTORi as a mechanistically rational strategy for reprogramming immune resistance in *TP53*-mutant HNSCC and provide a strong preclinical rationale for combining everolimus with immune therapy in patients who are likely to fail immunotherapy.

## Introduction

Head and neck squamous cell carcinoma (HNSCC) comprises a highly heterogeneous group of malignancies arising from the mucosal epithelium of the upper aerodigestive tract, including the oral cavity, pharynx, and larynx (1). It is the sixth most common cancer globally and accounts for an estimated 4.5% of newly diagnosed cancer cases and 4.6% of cancer-related deaths worldwide (2). Despite advances in locoregional management, outcomes for patients with advanced disease remain poor, underscoring the urgent need for more effective therapeutic strategies.

Comprehensive genomic profiling by The Cancer Genome Atlas (TCGA) has defined the mutational landscape of HNSCC and identified recurrent alterations in key oncogenic signaling networks driving tumor initiation and progression (3). Among these, *TP53* mutations represent the most frequent somatic genomic alterations in HNSCC, occurring in approximately 71.5% of all cases and in roughly 86% of HPV-negative tumors (3–7). The *TP53* gene produces the p53 protein, which safeguards genomic stability (8, 9). When mutated, this protective function is lost, leading to genomic instability that promotes cancer cell growth, invasion, and resistance to therapy (3, 4). Beyond its tumor cell-intrinsic effects, mutant p53 actively remodels the tumor microenvironment (TME) to promote immune evasion and therapeutic resistance (10). *TP53*-mutant tumors exhibit resistance not only to conventional chemo and radiotherapy, but also to a broad range of immune-based therapies, including immune checkpoint inhibitors (ICIs), chimeric antigen receptor (CAR) T cell therapy, and hematopoietic stem cell transplantation (HSCT) (10). Mechanistically, mutant p53 alters cytokine and chemokine profiles within the TME and reprograms myeloid populations toward immunosuppressive M2 macrophages and myeloid-derived suppressor cells (MDSCs), collectively suppressing effector T cell function and enabling tumor immune escape (11, 12). Compounding this, mutant p53 downregulates MHC class I and II expression while upregulating PD-L1, further disabling antitumor immune effector mechanisms and entrenching an immune-cold TME (10–14).

Importantly, loss of wild type p53 increases cellular reliance on the PI3K-Akt-mTOR pathway for survival and growth of tumor cells (6, 10, 15–17). This dependency on the PI3K-Akt-mTOR signaling axis is a critical and therapeutically exploitable consequence of *TP53* mutation. Wild type p53 normally restrains mTOR activity by driving transcription of key inhibitors, including tuberous sclerosis complex (TSC1/2) and Sestrin1/2 (18, 19). Loss of functional p53 dismantles these inhibitory checkpoints, resulting in persistent mTOR hyperactivation. Essentially, mutant p53 hijacks this pathway for oncogenic gain, inadvertently creating a system vulnerability that renders these tumors selectively dependent on, and thus susceptible to, mTOR inhibition (mTORi) (19, 20).

Our previous work demonstrated that mTOR inhibitors, including rapamycin and everolimus, significantly reduced tumor burden, suppressed angiogenesis, and prolonged survival in *TP53*-mutant HNSCC animal models with hyperactivated mTOR (21, 22). Everolimus treatment also increased MHC class I (MHC-I) expression on tumor cells, potentially enhancing tumor antigen presentation and immune recognition (23). Similarly, prior work in syngeneic HNSCC models demonstrated that rapamycin enhanced CD8+ T-cell expansion and, when combined with PD-L1 blockade, improved tumor control and prolonged survival in immunogenic tumors, thereby challenging the longstanding view of mTOR inhibitors as broadly immunosuppressive (24).

However, that study also revealed that poorly immunogenic tumors failed to respond to PD-L1 blockade, and whether mTORi could reprogram immune resistance in *TP53*-mutant, immune-cold, PD-1-refractory tumors remain unresolved. mTORi directly targets the nutrient-sensing and growth signaling pathways that cancer cells rely on for survival, thereby inhibiting tumor cell growth and proliferation (25). Beyond its direct effects on cancer cells, mTORi can reshape the tumor secretome by modulating protein synthesis and promoting autophagic degradation of secreted factors, with the potential to fundamentally alter immune cell recruitment and function within the TME (26–28). Therefore, we hypothesized that mTORi not only suppresses the tumor-intrinsic growth signaling but also modulates the TME to support effector immune function in *TP53*-mutant HNSCC. Specifically, we postulated that targeting mTOR would reverse mutant p53-driven immune-cold TME and create a more immune-permissive microenvironment.

In this study, we characterized the antitumor and immunomodulatory effects of the mTOR inhibitor everolimus in *TP53*-mutant, anti-PD-1 resistant HNSCC cell lines and syngeneic mouse models. Our study demonstrated that mTORi remodels the immune microenvironment by suppressing the HIF-1α/VEGF signaling, reducing PD-1/PD-L1 axis activation, and enhancing CD8+ T cell infiltration and function. Furthermore, we provide evidence that mTORi exerts both tumor-intrinsic and T cell-intrinsic effects. These findings support targeting mTOR as a rational strategy to enhance immunotherapeutic efficacy in *TP53*-mutant HNSCC.

## Materials and methods

### Cell lines and culture conditions

The *TP53*-mutant HNSCC cells lines ROC-1 and MOC-2 were used in this study. The ROC-1 cell line was generously provided by Dr. Jeffrey N. Myers, (MD Anderson Cancer Center, The University of Texas), and the MOC-2 cell line was provided by Dr. Clint T. Allen (National Institutes of Health, Bethesda). Cells were cultured according to the protocols established by the originating laboratories (29, 30). All in vitro experiments were performed using cells at 70 - 80% confluency and at passages below 20.

### Mice model and in vivo experiments

Female C57BL/6 mice, 8–10-weeks-old were procured from Jackson Laboratory and maintained under pathogen-free conditions. For tumor modeling, subconfluent ROC-1 (5×10^6^/injection) and MOC-2 (0.5×10^6^/injection) cells were mixed with matrigel matrix and injected subcutaneously into the flanks of the mice (29, 30). The tumors were allowed to reach 100-150 mm^3^ before initiating treatment. The tumor mice were randomly divided into vehicle control (n = 6) or treatment groups (n = 6). The control group received 1% carboxymethylcellulose sodium salt (CMC-Na) in water (oral gavage, once daily) and the treatment group received everolimus (RAD001; 5 mg/kg, oral gavage, once daily, Selleck Chemicals, S1120) suspended in 1% CMC-Na. Tumor dimensions were measured three times per week using fine digital calipers. Tumor volume was calculated using the formula: V (mm^3^) = length (mm) × width^2^ (mm)/2. The mice were considered to have reached study endpoints when tumor volume reached 1000 mm^3^ or when body weight decreased by more than 15%. For mechanistic analysis, tumors were harvested after 1 week of treatment with vehicle or everolimus. All procedures were conducted in accordance with the Institutional Animal Care and Use Committee (IACUC) guidelines at Louisiana State University Health Sciences Center, Shreveport (Protocol No.: P-25-037). Reagent details are provided in supplementary data Table S1.

### Flow cytometry with tumor tissues from mice

Freshly resected tumors were digested into a single cell suspension using the mouse tumor dissociation kit from Miltenyi Biotec following the manufacturer’s protocol. Flow cytometric analysis of tumor-infiltrating immune cell populations was performed using established protocols at the Center for Applied Immunology and Pathological Process (CAIPP) Immunophenotyping Core Facility at LSU Health Shreveport. Briefly, cells were first stained with zombie violet for live/dead discrimination and then treated with Fc block to prevent non-specific antibody binding. Cell surface staining was subsequently performed using the appropriate dilution of antibodies in FACS buffer.

For intracellular staining, cells were treated with 1X permeabilization buffer and then incubated with appropriate dilution of antibodies in 1X permeabilization. Afterwards, cells were washed with permeabilization/FACS buffer and fixed prior data acquisition on a flow cytometer.

Antibody panels, dilutions, and flow gating strategies are provided in the supplementary data Table S2 and Figure S3, S4, and S5.

### In vitro hypoxia exposure protocol

To investigate the effect of everolimus on hypoxia signaling, particularly the HIF-1α/VEGFA and HIF-1α/PD-L1 axes, we performed an in vitro hypoxia exposure experiment. Briefly, 0.4 × 10^6^ ROC-1 and MOC-2 cells were seeded in 60 mm culture dishes and incubated overnight in a CO_2_ incubator. The following morning, the culture media were replaced with fresh media containing 100 nM everolimus or vehicle (DMSO), and the culture dishes were placed in Coy’s hypoxia chamber maintained at 1% O_2_, 5% CO_2_, and 37°C for 5 hours. Protein samples were collected immediately after hypoxia exposure.

### Western blotting

Cultured cells and tumor tissues were lysed, respectively, in 1X cell lysis buffer, and 1X RIPA buffer supplemented with protease inhibitors, phosphatase inhibitors and 1mM PMSF. Cell lysis was assisted with sonication. After sonication, lysates were incubated on ice for 10 minutes and centrifuged at 14000 rpm for 15 minutes at 4 °C, and the supernatant was collected. The amount of protein in the collected supernatant was then quantified, mixed with Laemmli sample buffer, and heated at 96 °C for denaturation. The sample lysates containing equal amounts of protein (50 µg) were resolved by SDS-PAGE and transferred onto PVDF membranes using Bio-Rad Trans-Blot Turbo Transfer System. The membranes were then blocked in 3% bovine serum albumin (BSA) in TBST for 1.5 hours and incubated with 1:1000 dilutions of the primary antibodies overnight at 4 °C, followed by 1:1000 dilutions of secondary antibodies for 1.5 hours at room temperature (31). Protein bands were detected using a Bio-Rad ChemiDoc MP Imaging System. The antibodies and other reagents used are listed in supplementary data Table S1.

### Immunohistochemistry (IHC) for CD31

CD31 staining of tumor sections were performed following standard IHC protocol with slight modifications (31). Tumor tissues were harvested, fixed in 10% zinc buffered formalin, paraffin-embedded, and sectioned at 5 µm thickness. After deparaffinization and rehydration, antigen retrieval was performed in Tris buffer (pH 9.0) with boiling in a pressure cooker. Endogenous peroxidase activity was blocked using 3% hydrogen peroxide, and nonspecific binding was blocked with 5% goat serum in TBST. Sections were then incubated with an anti-CD31 primary antibody overnight at 4 °C. An HRP-conjugated secondary antibody was applied for 30 minutes, and staining was visualized using DAB chromogen.

Sections were counterstained with hematoxylin, dehydrated, mounted, and imaged by light microscopy. Vascular density was quantified based on CD31-positive staining in tumor sections using Fiji (ImageJ).

### Realtime-polymerase chain reaction (RT-PCR)

The expression level of various genes of interest were assessed by real-time PCR with slight modification to a standard protocol (31). In brief, total RNA was isolated from cells and tumor tissues using Trizol reagent, and cDNA was synthesized using iScript™ cDNA Synthesis Kit according to the kit protocol. Equal amounts of cDNA were amplified using gene-specific primers and SYBR green PCR Master Mix on a Bio-Rad CFX96 real-time PCR system. A standard thirty-five amplification cycle (95 °C for 10 s, 58 °C for 10 s, and 72 °C for 30 s) were performed. Gene expression levels were normalized to glyceraldehyde-3-phosphate dehydrogenase (GAPDH) using the ΔΔCT method. The sequences of the forward and reverse primers used are provided in supplementary data Table S6 and the reagents used are listed in supplementary data Table S1.

### Isolation of tumor-infiltrating CD8+ T Cells and cytotoxicity assay

Freshly resected tumors were dissociated into a single cell suspension using the mouse tumor dissociation kit per the manufacturer’s protocol. The obtained cell suspensions were then used to isolate CD8+ T cells. In brief, the cell suspensions were labelled with CD8 (TIL) MicroBeads and sorted in a QuadroMACS magnetic bead separation system using LS columns according to the manufacturer’s protocols.

Purity of the isolated CD8+ T cells was confirmed by flow cytometry (supplementary data Figure S7). Isolated CD8+ T cells from the tumors of vehicle and everolimus-treated mice were co-cultured with ROC-1 tumor cells prelabelled with Hoechst3342 (H33342) fluorescent dye at a 5:1 ratio. After 18 hours of incubation in a cell culture incubator, the co-cultures were centrifuged, and fluorescent intensity in the cell free supernatant was measured. T cell cytotoxicity kills the tumor cells, causing a proportionate release of H33342 into the media (32). Cytotoxicity was calculated relative to spontaneous and complete lysis controls using established formula. Spontaneous lysis was made with only H33342 labelled ROC-1 cells. Autolysis of cells releasing H33342 into the media was measured and defined as 0% cytotoxicity. Complete lysis was performed by treating H33342-labelled ROC-1 cells with a detergent (0.01% Triton X100). Detergent treatment lysed all ROC-1 cells, resulting in maximal release of H33342 into the media, which was defined as 100% cytotoxicity.

### In vitro assessment of everolimus impact on PD-1 expression and T cell function

PD-1+ (exhausted-like) T cells were generated with T cell receptor (TCR) stimulation of splenocytes (33 34). Briefly, RBC-lysed splenocytes from the naïve C57BL6 mice were cultured in RPMI1640 supplemented with 10% FBS and 1X penicillin-streptomycin and then stimulated on plates coated with anti-CD3ε (5 µg/mL) and anti-CD28 (1 µg/mL) antibodies. On day 3, cells were transferred to fresh media supplemented with 50 ng/mL mouse interleukin-2 (IL-2) with or without everolimus (100 nM) for an additional 3 days. On day 7, cells were harvested, stained, and analyzed for PD-1 expression by flow cytometry. Antibody panels, dilutions, and flow gating strategy are provided in the supplementary data Table S2 and Figure S8.

For cytotoxicity assays, CD8+ T cells were magnetically sorted from the cultured control and everolimus-treated splenocytes and then co-cultured with ROC-1 tumor cells as described above.

### Statistical analysis

Graphical representation and statistical analyses were performed using GraphPad prism v10. Data are presented as mean ± SEM. The tests of significance between the presented data are reported as *p*-values, with *p*-value < 0.05 considered statistically significant. Statistical significance between two groups was determined using an unpaired two-tailed Student’s T-test. Comparisons among multiple groups were performed using one-way analysis of variance (ANOVA) followed by Tukey’s multiple comparisons. Error bars reflect the standard error of means (SEM). Significance levels are indicated as follows: nonsignificant (ns) (*p* ≥ 0.05); * (*p* < 0.05); ** (*p* < 0.01); *** (*p* < 0.001).

## Results

### mTOR inhibition alters the immune-cold TME promoting immune cell infiltration in *TP53*-mutant HNSCC

Mutant p53 induces immune-cold TME and promotes therapeutic resistance in HNSCC (29, 30). Loss of p53 function in HNSCC increases cellular reliance on the mTOR pathway for survival and growth, resulting in constitutive activation of mTOR (15–19). Given this dependency, we investigated whether pharmacologic mTORi could suppress tumor growth and remodel the immune-cold TME in *TP53*-mutant HNSCC.

In this study, treatment with the mTOR inhibitor everolimus significantly suppressed the growth of *TP53*-mutant, immune-cold ROC-1, and MOC-2 syngeneic tumor grafts (Fig 1A). Next, to assess the effects of mTORi on the immune contexture of these tumors, we subjected tumor tissues to multiparameter flow cytometry (Fig. 1B). Flow cytometric analysis revealed a significant increase in overall immune cell infiltration in everolimus-treated tumors, with notable increases in CD8+ T cells and dendritic cells (DCs) (Fig. 1C, 1D). Everolimus treatment also significantly reduced the infiltration of immunosuppressive regulatory T cells (Tregs), and PD-1 expressing (exhausted-like) CD4+ and CD8+ T cells. However, no significant changes were observed in natural killer (NK) cells or macrophage infiltration.

**Figure 1.**
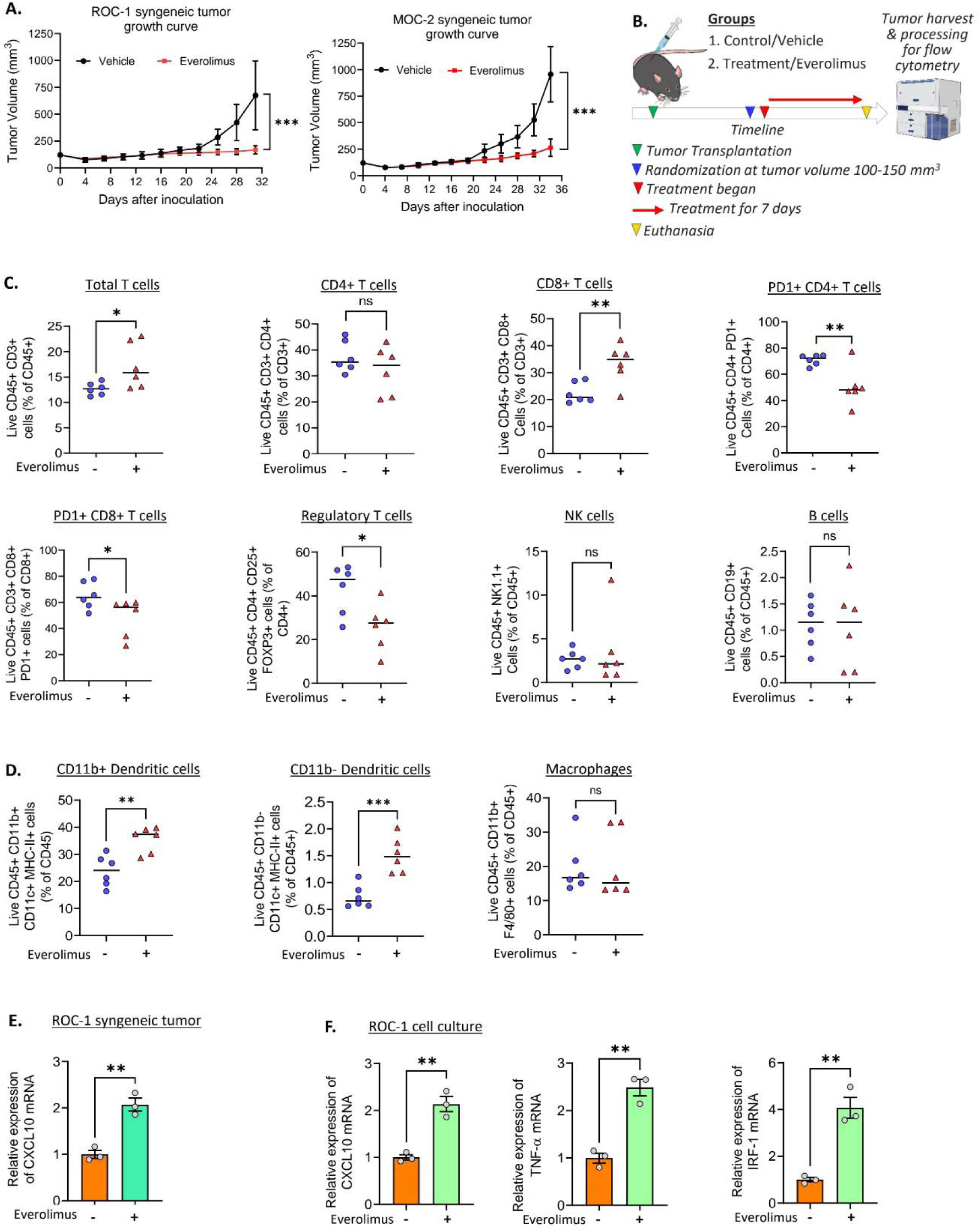
mTOR inhibition with everolimus reshapes *TP53*-mutant immune-cold TME. (A) Growth curves of syngeneic ROC-1 and MOC-2 tumors showing significant suppression of tumor growth following daily oral administration of everolimus (5 mg/kg body weight). Data are presented as mean ± SEM, n = 6 mice/group. (B) Schematic overview of the treatment and tumor harvest timeline for immune phenotyping of ROC-1 syngeneic tumors by flow cytometry. (C) Flow cytometric characterization of lymphocytic immune infiltration into vehicle and everolimus-treated ROC-1 syngeneic tumor, showing data on total immune infiltration with CD3+ cells, CD4+ T cell, CD8+ T cell, PD1+ (exhausted-like) CD4+ and CD8+ T cells, regulatory T cells (Tregs), B lymphocytes, and NK cells. Six individual tumors from each condition were analyzed. (D) Flow cytometric characterization of myeloid immune infiltration into vehicle and everolimus-treated ROC-1 syngeneic tumor, showing data for CD11b+ and CD11b-dendritic cells (DCs), as well as total macrophages (CD11b+ F4/80+ cells). Six individual tumors from each condition were analyzed. (E) RT-PCR analysis showing CXCL10 mRNA expression in tumor tissues. Data are presented as mean ± SEM, n = 3 (RNA was extracted from 3 independent experiments). (F) RT-PCR analysis showing CXCL10, TNF-α, and IRF-1 mRNA expression in ROC-1 cells with vehicle and everolimus treatment. Data are presented as mean ± SEM from n = 3 independent experiments. Statistical significance between two groups was determined using an unpaired two-tailed Student’s T-test. Significance levels are indicated as follows: nonsignificant (ns) (p ≥ 0.05); * (p < 0.05); ** (p < 0.01); *** (p < 0.001).

To evaluate immune-related transcriptional changes in everolimus-treated tumors, we performed RT-PCR on tumor-derived RNA. Based on our flow cytometry results showing significantly increased infiltration of CD8+ effector T cells, we examined the mRNA expression of C-X-C motif chemokine ligand 10 (CXCL10) in tumor tissues. CXCL10 is a major mediator of CD8+ T-cell recruitment into tumors and promotes anti-tumor immunity in solid tumor models (35).

RT-PCR analysis of cDNA synthesized from tumor-derived RNA revealed significantly higher CXCL10 mRNA levels in everolimus-treated tumors (Fig. 1E). Consistent with this, in vitro treatment of ROC-1 cells with everolimus increased CXCL10 mRNA expression, possibly through an interferon regulatory factor-1 (IRF-1)-regulated axis (Fig. 1F) (36). Additionally, everolimus increased expression of the pro-inflammatory cytokine TNF-α in ROC-1 cells, which is known to promote inflammation, immune cell recruitment, and immune response (37). TNF-α has also been reported to increase CXCL10 expression (38). These findings suggest that tumor cells may represent one source of CXCL10 within the tumor microenvironment, in addition to stromal and immune components. Together, these results indicate that mTORi alters cytokine and chemokine signaling to regulate immune cell infiltration dynamics in a mutant *TP53*-driven, immune-cold TME.

### mTOR inhibition suppresses the HIF-1α/VEGFA angiogenic axis and reduces MDSC recruitment

mTOR signaling plays crucial role in regulating HIF-1α and VEGFA activity (39, 40). mTOR regulates cap-dependent translation, rendering short-lived proteins like HIF-1α highly sensitive to its inhibition. This sensitivity arises from a dual mechanism: the suppression of protein synthesis and the simultaneous activation of autophagy-lysosomal and UPS-mediated degradation (41, 42). Furthermore, as HIF-1α transcriptionally regulates and promotes VEGFA protein expression, which in turn promotes immune evasion by recruiting MDSCs (43), we investigated whether mTORi disrupts the HIF-1α/VEGFA axis in *TP53*-mutant HNSCC. To confirm target engagement and the efficacy of mTORi with everolimus at the concentration used, we assessed pS6 levels in treated tumor cells in vitro and in tumors in vivo and observed a significant reduction in pS6 protein by western blot. Consistent with effective pathway inhibition, treatment of ROC-1 and MOC-2 cell cultures with everolimus significantly reduced basal HIF-1α and VEGFA protein levels (Fig. 2A). In line with these in vitro findings, western blotting of tumor lysates from everolimus-treated syngeneic mice revealed significantly reduced HIF-1α and VEGFA protein levels in vivo (Fig. 2B). To determine whether the reduced VEGFA levels translated into functional inhibition of tumor angiogenesis in vivo, we performed CD31 IHC staining of tumor sections. CD31 IHC revealed significant inhibition of angiogenesis in everolimus-treated tumors, as demonstrated by reduced vascular density (Fig. 2C). Consistent with the observed reduction in VEGFA, a key driver of MDSC recruitment and differentiation within the TME (43), flow cytometric analysis of tumor tissues revealed significantly reduced frequencies of monocytic and granulocytic MDSCs in everolimus-treated tumors (Fig. 2D). Together, these findings demonstrate that mTORi suppresses the HIF-1α/VEGFA angiogenic axis, thereby limiting the recruitment of immunosuppressive myeloid cells.

**Figure 2:**
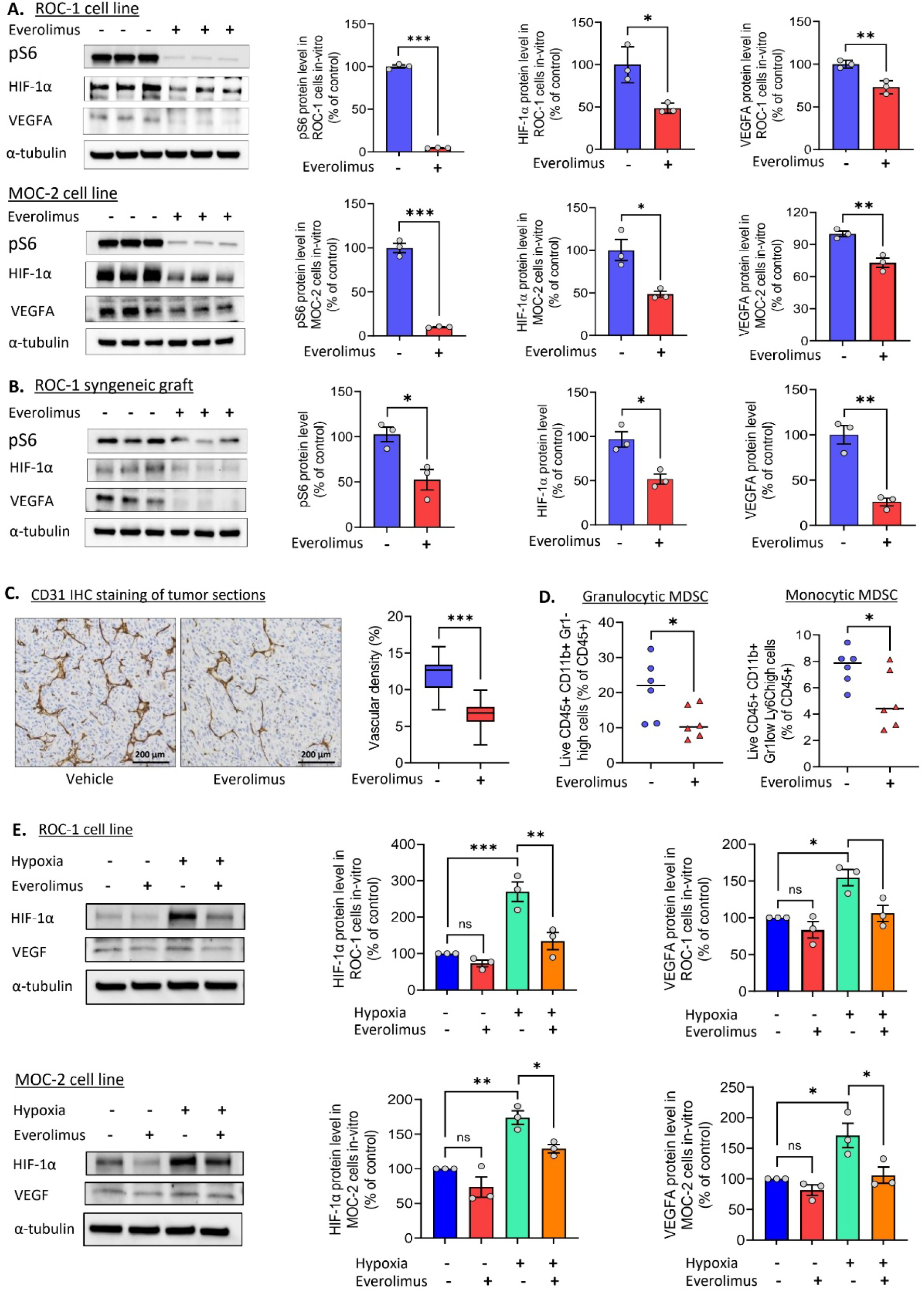
mTOR inhibition alters the HIF-1α/VEGFA signaling in *TP53*-mutant HNSCC and affects MDSC recruitment. (A) Western blot analysis of HIF-1α and VEGFA protein levels in cultured ROC-1 and MOC-2 cells treated with everolimus (100 nM for 24 hours). Representative immunoblots and densitometry plots were shown. Data are presented as Mean ± SEM from n = 3 independent experiments. (B) Western blot analysis of syngeneic ROC-1 tumor lysate from vehicle (1% CMC-Na in water) and everolimus-treated mice (5 mg/kg body weight daily). Representative immunoblots and densitometry plots were shown. Data are presented as Mean ± SEM from n = 3 independent animal experiments. (C) Representative CD31 immunohistochemical staining of tumor sections showing reduced angiogenesis in everolimus-treated tumors. A box-and-whisker plot shows a difference in vascular density (%) between vehicle and everolimus-treated tumors. Sections from six individual tumors were stained and analyzed. (D) Granulocytic and monocytic MDSCs in the control and everolimus-treated syngeneic ROC-1 tumors. Six individual tumors from each condition were analyzed. (E) Hypoxia-induced HIF-1α /VEGFA signaling and effect of everolimus. ROC-1 and MOC-2 cell cultures were exposed to hypoxia (1% 0_2_, 5% CO_2_, 37 °C temperature) with and without everolimus (100 nM in DMSO). Proteins were harvested immediately after 5 hours. Representative immunoblots and densitometry plots are shown. Data represents mean ± SEM from n = 3 independent experiments. Statistical significance between two groups was determined using an unpaired two-tailed Student’s T-test. Comparisons among multiple groups were performed using one-way analysis of variance (ANOVA) followed by Tukey’s multiple comparisons. Significance levels are indicated as follows: nonsignificant (ns) (p ≥ 0.05); * (p < 0.05); ** (p < 0.01); *** (p < 0.001).

Hypoxia is a major physiological inducer of HIF-1α in the TME. We therefore examined whether everolimus could suppress hypoxia-driven HIF-1α signaling. In hypoxia exposure experiments, pretreatment of ROC-1 and MOC-2 cells with everolimus significantly prevented hypoxia-induced increases in both HIF-1α and VEGFA protein levels (Fig. 2E). This further confirms that mTORi disrupts hypoxia-dependent angiogenic signaling.

### mTOR inhibition reduces PD-L1 expression through Myc-STAT3 and hypoxia-dependent mechanisms

Mutant p53 is well established to increase PD-L1 expression and immune evasion in cancer (44). Given the over-reliance of *TP53*-mutant tumors on mTOR (6, 10, 15–17), we examined whether mTORi modulates mutant p53-induced PD-L1 expression. In vitro treatment of ROC-1 and MOC-2 cell cultures with everolimus significantly reduced PD-L1 protein levels, as demonstrated by western blot (Fig 3A). Because HIF-1α is a major transcriptional regulator of PD-L1 in tumors, this reduction in PD-L1 could partly be due to previously observed suppression of HIF-1α by everolimus (Fig 2A, 2B). We next examined the other key PD-L1 transcriptional regulators c-Myc and pSTAT3, both of which bind the PD-L1 promoter to drive its expression (45). Everolimus treatment significantly reduced c-Myc and pSTAT3 levels in both cell lines, implicating transcriptional suppression as a contributing mechanism. Consistent with these in vitro findings, everolimus-treated syngeneic tumors showed significantly reduced c-Myc, pSTAT3, and PD-L1 protein levels in vivo (Fig. 3B).

**Figure 3:**
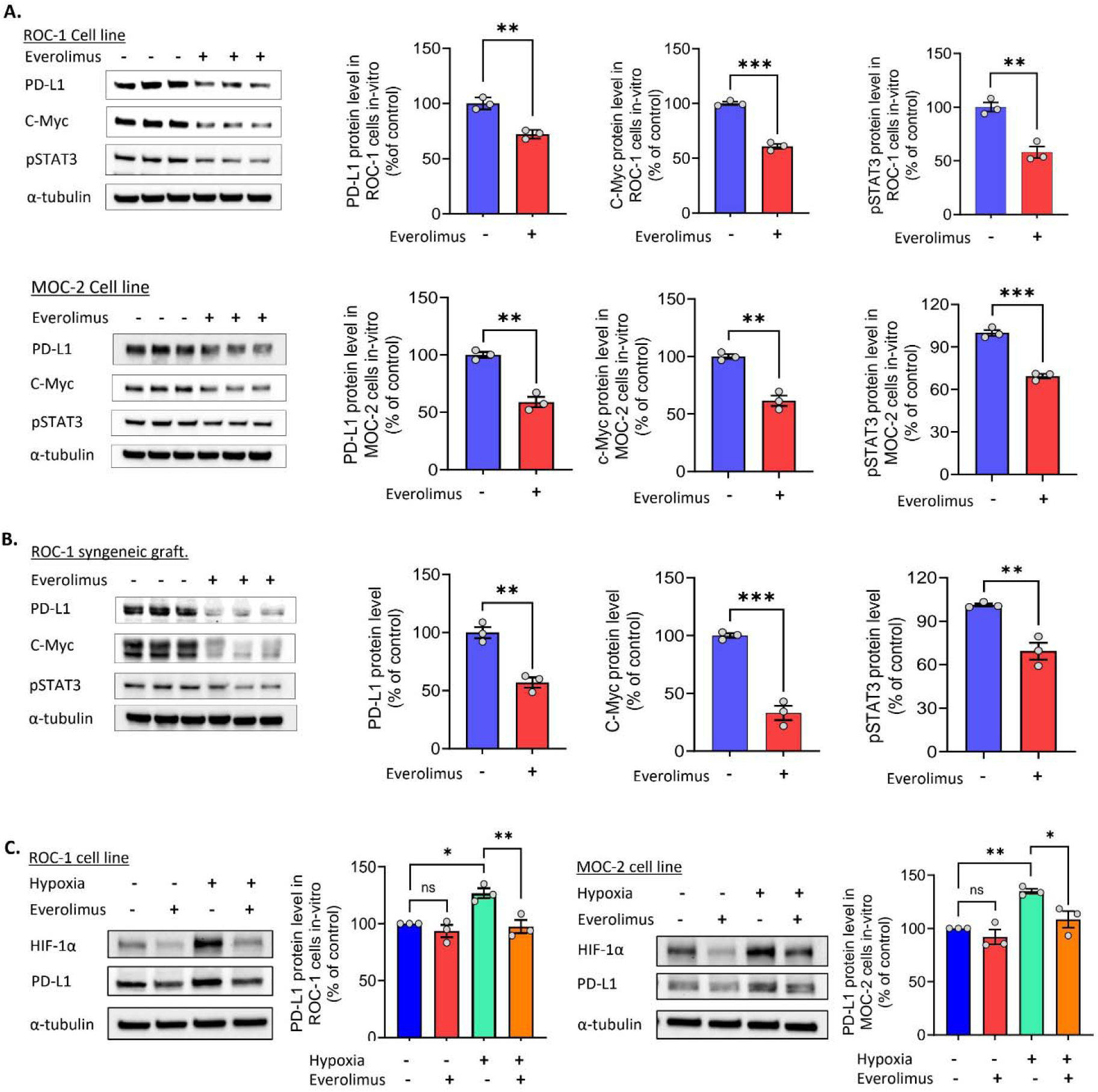
mTOR inhibition regulates PD-L1 expression in *TP53*-mutant HNSCC through transcriptional and hypoxia dependent mechanisms. (A) Western blot analysis of PD-L1 protein and its transcriptional regulator c-Myc and pSTAT3 levels in cultured ROC-1 and MOC-2 cells treated with everolimus (100 nM for 24 hours). Representative immunoblots and densitometry plots were shown. Data are presented as mean ± SEM from n = 3 independent experiments. (B) Western blot analysis with syngeneic ROC-1 tumor lysate from vehicle (1% CMC-Na in water) and everolimus treated mice (5 mg/kg body weight daily) for PD-L1, c-Myc, and pSTAT3 protein levels. Representative immunoblots and densitometry plots were shown. Data are presented as mean ± SEM from n = 3 independent animal experiments. (C) Hypoxia-induced HIF-1α signaling in PD-L1 expression and effect of everolimus. ROC-1 and MOC-2 cell cultures were exposed to hypoxia (1% 0_2_, 5% CO_2_, 37 °C temperature) with and without everolimus (100 nM in DMSO). Proteins were harvested immediately after 5 hours. Representative immunoblots and densitometry plots were shown. Data are presented as mean ± SEM from n = 3 independent experiments. Statistical significance between two groups was determined using an unpaired two-tailed Student’s T-test. Comparisons among multiple groups were performed using one-way analysis of variance (ANOVA) followed by Tukey’s multiple comparisons. Significance levels are indicated as follows: nonsignificant (ns) (p ≥ 0.05); * (p < 0.05); ** (p < 0.01); *** (p < 0.001).

Since hypoxic TME induces PD-L1 expression in tumor cells (46), we next evaluated whether mTORi affects hypoxia-driven PD-L1 expression. ROC-1 and MOC-2 cells exposed to hypoxic conditions (1% O_2_) demonstrated robust induction of both HIF-1α and PD-L1. However, everolimus pretreatment prevented hypoxia-induced accumulation of HIF-1α and PD-L1 proteins (Fig. 3C). These results indicate that mTORi suppresses PD-L1 expression through convergent mechanisms, attenuating c-Myc/STAT3-driven transcription and blocking hypoxia-dependent induction.

### mTOR inhibition therapy enhances the cytotoxic competence of tumor-resident CD8+ T cells

Treatment with everolimus attenuated Tregs, MDSCs infiltration, and tumor cells’ expression of PD-L1 while also expanding CD8+ T cell and DC infiltration into the tumors. Given that these changes can potentially impact effector T cell function, we assessed the cytotoxicity of CD8+ T cells (Fig. 4A). Ex vivo cytotoxicity assays showed that CD8+ T cells isolated from everolimus-treated ROC-1 syngeneic tumors exhibited a significantly greater ability to kill ROC-1 tumor cells compared with CD8+ T cells from vehicle-treated control tumors (Fig. 4B). Consistent with this, intracellular flow cytometry revealed higher IFN-γ expression in CD8+ T cells from everolimus-treated tumors (Fig. 4C). RT-PCR analysis of RNA from CD8+ T cells isolated from everolimus-treated tumors showed significantly elevated granzyme B mRNA levels compared to controls (Fig. 4D). Everolimus-treated tumors also expressed significantly lower T cell immunoreceptor with Ig and ITIM domains (TIGIT) mRNA levels, an inhibitory receptor known to suppress T-cell effector function, indicating enhanced immune competency of tumor-infiltrating T cells (Fig. 4E).

**Figure 4:**
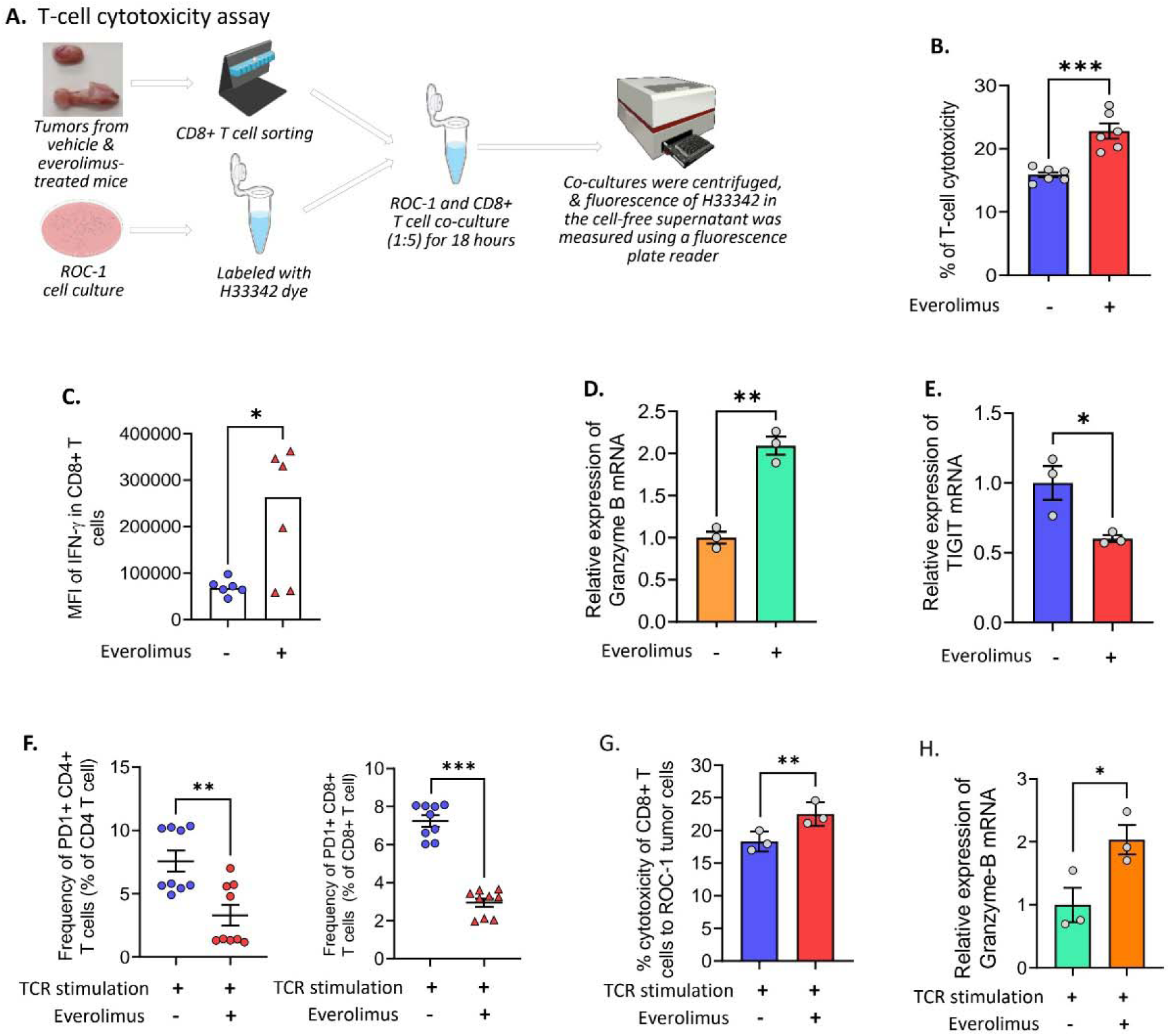
mTOR inhibition with everolimus enhances effector (CD8+) T cell function and attenuates PD-1 expression. (A) Schematic overview of the in vitro T cell cytotoxicity assay using tumor-infiltrating CD8+ T cells. (B) Representative bar diagram showing T cell cytotoxicity (%) using CD8+ T cells isolated from tumors of vehicle- and everolimus-treated mice. Data represents mean ± SEM of six independent observations, n = 6. (C) Flow cytometric analysis showing the mean fluorescence intensity (MFI) of IFN-γ in CD8+ T cells from control and everolimus-treated ROC-1 syngeneic tumors. Surface and intracellular flow staining was performed with the antibodies for required markers. Six individual tumors from each condition were analyzed. (D) RT-PCR analysis showing mRNA levels of cytotoxic mediator granzyme-B in tumor infiltrating CD8+ T cells. Data are presented as mean ± SEM. n = 3 (RNA was extracted from T cells isolated from pulled tumors separately from 3 independent experiments). (E). RT-PCR analysis showing TIGIT mRNA expression in tumor infiltrating CD8+ T cells. Data are presented as mean ± SEM. n = 3 (RNA was extracted from T cells isolated from pulled tumors separately from three independent experiments). (F) Flow cytometric analysis showing the effect of mTOR inhibition on PD-1 expression in CD4+ and CD8+ T cells following sustained TCR-stimulation of splenocytes with anti-CD3ε and anti-CD-28 antibodies. Data are presented as mean ± SEM from n = 3 independent experiments in triplicate. (G) Representative bar diagram showing T cell cytotoxicity (%) using CD8+ T cells isolated from control and everolimus-treated splenic cells following TCR-stimulation for PD-1 expression. ROC-1 cells were used as Target. Data are presented as mean ± SEM from n = 3 independent experiments. (H) RT-PCR showing mRNA levels of cytotoxic mediator granzyme-B in CD8+ T cells isolated from control and everolimus-treated splenic cells following TCR stimulation. Data are presented as mean ± SEM from n = 3 independent experiments. Statistical significance between two groups was determined using an unpaired two-tailed Student’s T-test. Significance levels are indicated as follows: nonsignificant (ns) (p ≥ 0.05); * (p < 0.05); ** (p < 0.01); *** (p < 0.001).

T-cell exhaustion, characterized by PD-1 expression, significantly impairs T cell function by reducing proliferation, cytokine production, and killing capacity (47). Our flow cytometric analysis of tumor tissues revealed significantly reduced frequencies of PD-1+ (exhausted-like) CD4+ and CD8+ T cells in everolimus-treated tumors (Fig. 1C). We therefore investigated whether mTORi directly modulates T cell-intrinsic PD-1 expression. We employed an in vitro exhaustion model using sustained TCR stimulation with anti-CD3ε and CD28 antibodies to drive PD-1 induction. Everolimus treatment significantly reduced frequencies of PD-1+ (exhausted-like) CD4+ and CD8+ T cells following sustained TCR stimulation (Fig. 4F). Concurrently, the CD8+ T cells isolated from everolimus-treated cultures exhibited significantly enhanced cytotoxicity against ROC-1 tumor cells (Fig. 4G), with RT-PCR confirming elevated granzyme B expression (Fig. 4H). These findings indicate that mTORi attenuates PD-1-driven T cell exhaustion and restores effector cytotoxic function through direct T cell-intrinsic mechanisms, providing a basis for its activity in *TP53*-mutant, PD-1-resistant HNSCC.

## Discussion

The PI3K/AKT/mTOR signaling pathway is frequently hyperactivated in *TP53*-mutant HNSCC, creating a tumor cell dependency that renders this disease subset selectively vulnerable to mTOR inhibitors (4, 20). While mTOR inhibitors have traditionally been considered immunosuppressive (48), emerging evidence indicates that they can modulate immune responses (49). The immunological consequences of mTORi, however, appear to depend on treatment context, including dosing, timing, and the type of specific inhibitor used (48, 50). More recent studies have identified immunostimulatory effects of mTORi, particularly in promoting memory CD8+ T cell generation (51). In the present study, we characterized the immunostimulatory effects of everolimus in *TP53*-mutant, PD-1-resistant HNSCC and provide evidence that mTORi augments T cell function by reshaping the TME and attenuating PD-1/PD-L1 checkpoint engagement. We utilized the *TP53*-mutant HNSCC cell lines ROC-1 and MOC-2, as well as their corresponding syngeneic flank tumor models, which are all well-established for recapitulating immune-cold TMEs resistant to immunotherapy (29, 30, 52). *TP53* mutations represent an early and frequent event in human HNSCC development, and these models closely mimic the aggressive nature of human disease. Previous studies have demonstrated that ROC-1 and MOC-2 tumors are highly tumorigenic and display aggressive biological behavior (29, 30, 52). ROC-1 tumors in particular exhibit high lymphatic metastasis similar to human HNSCC and display an immune-cold TME (29). A similar immune-cold phenotype has been reported for MOC-2 tumors (30, 52). These models therefore provided an appropriate and clinically relevant platform to interrogate whether mTORi could remodel the immune-cold, immunotherapy-resistant TME driven by mutant p53.

Our findings demonstrate that mTORi profoundly alters the intratumoral immune cell infiltration, thereby modulating the TME. Mechanistically, everolimus increased expression of pro-inflammatory factors TNF-α and CXCL10 in ROC-1 cells, suggesting induction of a pro-inflammatory state with mTORi (37). CXCL10 is associated with recruitment of CD8+ effector T cells (35, 53). Consistent with this transcriptional change, flow cytometric analysis of tumor tissues revealed increased infiltration of CD8+ T cells, indicating a shift toward a more immune-permissive TME. Additionally, everolimus increased IRF-1 expression in ROC-1 cells. Prior studies have identified IRF-1 and TNF-α as inducers of CXCL10; consistently, the increased CXCL10 mRNA expression observed in ROC-1 cells may be mediated, at least in part, through an IRF-1 and TNFα-associated mechanism (36, 38). These results also suggest that tumor cells represent one source of CXCL10, although stromal and immune cells are also likely contributors. Importantly, our flow analysis of tumor tissues revealed significant increase in DCs following everolimus treatment. Given the central role of DCs in antigen presentation and T-cell priming (54), increased DC infiltration further supports the possibility that mTORi enhances immune activation within this p53-driven, immune-cold tumor model.

Given that immune cell function is strongly influenced by the hypoxic features of the TME, we examined how mTORi affects hypoxia-associated pathways. HIF-1α and VEGFA signaling play central roles in shaping immunosuppressive TME by promoting angiogenesis and immune exclusion (43, 55). Wild type p53 represses HIF-1α activity, whereas mutant p53 promotes HIF-1α and VEGFA expression, fostering angiogenesis and immunosuppression (55, 56). HIF-1α activity is further regulated by an mTOR signaling motif and STAT-3-dependent transcription and translational control via 4E-BP1 and S6K1 (39, 40). In addition, mTORi potentially lowers HIF-1α levels by suppressing protein synthesis and inducing autophagy-lysosomal degradation and UPS activity (41, 42). Consistent with this model of mTOR-dependent HIF-1α regulation, our analysis revealed significant downregulation of HIF-1α/VEGFA angiogenic signaling with mTORi. Western blot analysis confirmed reduced HIF-1α and VEGFA protein levels in ROC-1 and MOC-2 cell cultures and ROC-1 syngeneic tumor following everolimus-treatment. These results align with prior reports implicating mTORC1 in regulating HIF-1α accumulation through both transcriptional and translational mechanisms (22, 39–42). HIF-1α and VEGFA profoundly influence immune composition within the TME (43, 55). HIF-1α-driven VEGFA promotes abnormal vasculature, impairs immune cell trafficking, drives MDSC recruitment and expansion, and supports Treg accumulation within the TME (24, 55). In our study, flow cytometric analysis of tumors demonstrated significant reductions in MDSC subsets following mTORi, consistent with prior studies in oral cancer models treated with mTOR inhibitors (30). MDSCs and Tregs suppress antitumor immunity by producing tolerogenic cytokines such as IL-10 and TGF-β (57). Reduced infiltration of these suppressive populations therefore indicates that mTORi reprograms the TME toward an immune-permissive state.

Our study systematically characterized NK cell populations within the TME following everolimus treatment and found no significant changes in NK cell infiltration. In the PD-1-resistant setting, NK cells represent an important alternative cytotoxic effector population (58), and mTORi has been shown to modulate NK cell function (59). Notably, prior work in syngeneic oral cancer models demonstrated that the antitumor activity of rapamycin combined with PD-L1 blockade was CD8+ T cell-dependent and NK cell-independent, suggesting that NK cell contributions may be limited in this disease setting (24). The observed results suggest a similar mechanism operates in *TP53*-mutant, PD-1-resistant models.

The PD-1/PD-L1 axis represents a central mechanism of immune evasion in both HPV-positive and HPV-negative *TP53*-mutant HNSCC. Elevated PD-L1 expression in HNSCC and other cancer types exhibiting *TP53* mutations are associated with immune evasion and tumor progression (60–62). Prior studies have reported paradoxical PD-L1 upregulation following mTORi in certain cancer types (63, 64). In contrast, our study demonstrated significant reduction in PD-L1 expression with mTORi in both cultured HNSCC cells and syngeneic tumor models.

This discrepancy likely reflects a tumor-specific regulatory mechanism. In HNSCC, mutant p53 cooperates with its cofactor YAP to form a complex that enhances c-Myc stability and its binding to target promoters including PD-L1, thereby driving PD-L1 transcription (65, 66). On the other hand, mTOR promotes c-Myc translation and stability, which is required for PD-L1 expressions, and mTORi therefore may reduce PD-L1 expression indirectly through c-Myc suppression (67). Our data supports this mechanism, as reduced PD-L1 levels correlated with decreased c-Myc levels in both cell lines and syngeneic tumors. HIF-1α has also been shown to directly regulate PD-L1 transcription by binding to its promoter under hypoxic conditions (68). Consistent with this, in our study, hypoxic exposure of ROC-1 and MOC-2 cells increased PD-L1 expression, whereas everolimus prevented hypoxia-induced PD-L1 upregulation by suppressing HIF-1α accumulation. Collectively, these findings demonstrate that mTORi limits immunosuppression by targeting both oncogenic and hypoxia-driven PD-L1 regulations. Prior studies have shown blockade of HIF-1α-driven PD-L1 signaling reduces MDSCs’ production of IL-6 and IL-10, thereby enhancing T-cell function (68). Our findings that everolimus suppresses both HIF-1α and PD-L1 support the existence of a PD-L1/MDSC/T cell regulatory axis in *TP53*-mutant HNSCC.

Mutant p53 promotes immune evasion by impairing antigen presentation, enhancing PD-L1 expression, recruiting suppressive immune populations, and secreting immunomodulatory factors that inhibit T cell function (10–14). Our data indicate that mTORi reverses this phenotype by reducing Treg and MDSC infiltration and lowering tumor PD-L1 expression. Functionally, these changes translated into improved T cell effector activity. CD8+ T cells isolated from everolimus-treated tumors exhibited significantly enhanced cytotoxicity compared with controls. Gene expression analysis revealed elevated mRNA levels of granzyme B, consistent with enhanced cytotoxic function.

PD-1 is a key inhibitory receptor that promotes T cell exhaustion under chronic antigen exposure (47). Persistent PD-1 signaling diminishes T cell cytotoxicity and disrupts their metabolic process by suppressing glycolysis while promoting fatty acid oxidation, thereby cutting down the energy that is vital for active immune responses (47). In our study, multiparameter flow cytometry revealed a significantly lower percentage of PD-1-expressing CD4+ and CD8+ T cells in everolimus-treated tumors. This observation aligns with emerging evidence that mTORi can modulate PD-1 expression and promote less exhausted T cell phenotypes, thereby restoring their effector function (69–71). The combined reduction of PD-L1 on tumor cells and PD-1 on T cells indicates that mTORi disrupts inhibitory checkpoint signaling at both sides of the immunological synapse within the TME. RT-PCR analysis further revealed reduced TIGIT expression in tumor-infiltrating CD8+ T cells from everolimus-treated mice, providing additional evidence of enhanced functional competence (72). To validate these findings mechanistically, we performed in vitro T cell exhaustion assays. Chronic TCR stimulation induced PD-1 expression in T cells, whereas everolimus significantly reduced PD-1 induction and improved T cell cytotoxicity in co-culture assays. These findings were consistent with our in vivo observations, confirming a direct T cell-intrinsic effect of mTORi.

Prior clinical trials of single agent mTOR inhibitors, including temsirolimus, in recurrent and metastatic HNSCC demonstrated modest antitumor activity, suggesting that mTORi alone is insufficient for durable disease control in this setting (73). Our mechanistic findings provide a potential explanation for these clinical observations. Single-agent mTORi may prime the TME for immune attack without fully overcoming the depth of immune exclusion in *TP53*-mutant disease. This reframes the clinical opportunity: rather than pursuing mTOR inhibitors as standalone agents, our data support their use as TME-priming agents in combination with anti-PD-1 therapy. Critically, our study was conducted in a PD-1-resistant syngeneic model, and while we demonstrate that everolimus remodels the immune-cold TME toward an immune-permissive state, we did not directly test the everolimus and anti-PD-1 combination in vivo. This represents the next experimental step, and our mechanistic data now provide the preclinical rationale to pursue this combination specifically in *TP53*-mutant, PD-1-refractory HNSCC, the patient population with the greatest unmet clinical need.

Taken together, our study reveals that mTORi orchestrates a coordinated, multidimensional reprogramming of the mutant p53-driven immune-cold TME in HNSCC. As summarized in the proposed working model (Fig. 6), everolimus suppresses the HIF-1α/VEGFA angiogenic axis, reducing MDSC recruitment and relieving a major source of immune exclusion. Concurrently, mTORi attenuates PD-L1 expression through convergent c-Myc/STAT3-driven transcriptional suppression and blockade of hypoxia-dependent induction, dismantling checkpoint engagement at the tumor cell surface. These tumor-extrinsic changes are compounded by direct T cell-intrinsic effects, as mTORi reduces PD-1 expression while restoring cytotoxic effector function in CD8+ T cells. The net result is a shift from an immune-cold, immunotherapy-resistant TME toward an immune-permissive one characterized by robust DC infiltration, effector CD8+ T cell expansion, and reduced suppressive immune populations. Given the high prevalence of *TP53* mutations in HNSCC, the clinical availability of everolimus, and the urgent unmet need in PD-1-refractory disease, these findings provide a compelling mechanistic and translational rationale for combining mTORi with anti-PD-1 therapy in *TP53*-mutant, immunotherapy-resistant HNSCC. Clinical evaluation of this combination in this molecularly defined patient population is warranted.

**Figure 5:**
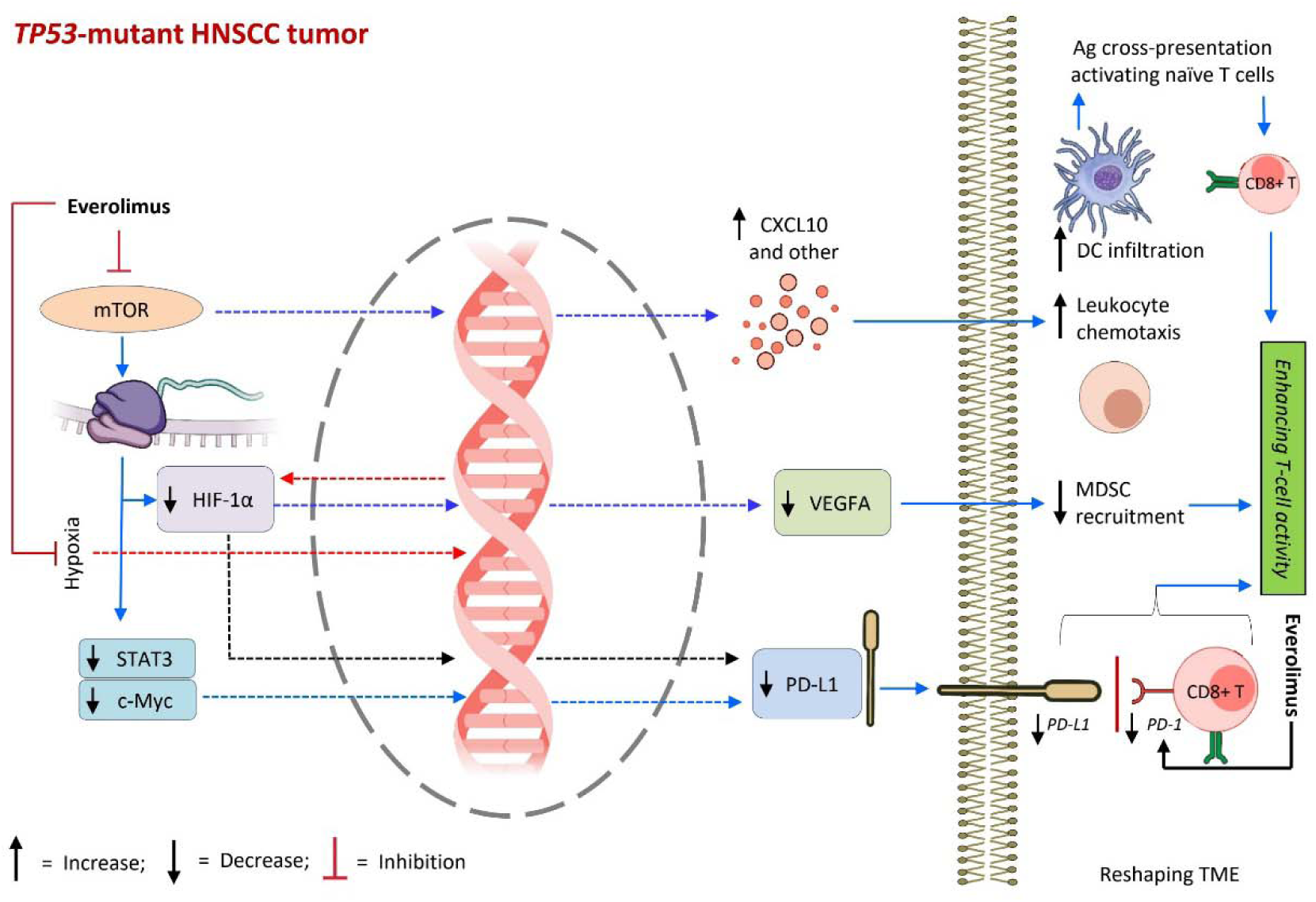
A summary figure showing a coordinated, multidimensional reprogramming of the mutant p53-driven immune-cold TME in HNSCC with everolimus.

## List of Abbreviations

mTOR: mechanistic target of rapamycin
mTORi: inhibition of mechanistic target of rapamycin
HNSCC: head and neck squamous cell carcinoma
PD-1: programmed cell death protein 1
PD-L1: programmed death-ligand 1
TME: tumor microenvironment
DCs: dendritic cells
HIF-1α: hypoxia-inducible factor-1 alpha
VEGFA: vascular endothelial growth factor A
MDSCs: myeloid-derived suppressor cells
ICI: immune checkpoint inhibitor
CXCL10: C-X-C motif chemokine ligand 10
TNF-α: tumor necrosis factor-alpha
IRF-1: interferon regulatory factor 1
TIGIT: T cell immunoreceptor with Ig and ITIM domains
TCR: T cell receptor

## Data Availability

All data described in this publication are available from the corresponding author, [C.O. Nathan], upon reasonable request.

## Acknowledgements

The authors sincerely acknowledge the research core facilities at LSU Health, Shreveport, including the Center of Applied Immunology and Pathological Processes Immunophenotyping Core supported by the NIH/NIGMS CoBRE Award P20 GM134974 (RRID:SCR_024781) for assisting in flow cytometry and Redox Molecular Signaling Core (RRID:SCR_024778) for the Coy hypoxic chamber.

## Author Contributions

**PN:** Conceptualization, Data curation, Formal analysis, Investigation, Methodology, Validation, Visualization, Writing-original draft, Writing-review and editing, **AK:** Conceptualization, Methodology, Writing-review and editing, **CL:** Methodology, **TMM:** Methodology, Resources, **SSV:** Methodology, **VAA:** Methodology, **OFC:** Resources, Writing-review and editing, **JSG:** Resources, Writing-review and editing, **CON:** Conceptualization, Funding acquisition, Supervision, Resources, Writing-review and editing.

## Consent for Publication

All authors have reviewed the article and have provided consent for publication.

