## Supplemental Table 1 for "mTOR inhibition augments antitumor immune effector response by reprogramming the *TP53*-mutant, immune-cold HNSCC tumor microenvironment"

**Supplementary Table- S1.**

| **Cell Culture Reagents** | **Manufacturer** | **Catalog #** |
| --- | --- | --- |
| DMEM, 1X | Corning | 10-017-CV |
| RPMI1640 | Cytiva/Hyclone | SH30096.01 |
| IMDM Modified | Cytiva/Hyclone | SH30228.02 |
| Nutrient Mixture | Cytiva/Hyclone | SH30026.01 |
| Setal Bovine Serum (FBS) | R&D Systems | S11150H |
| MEM NEAA (100X) | Gibco | 11140-050 |
| Sodium Pyruvate (100 mM) | Gibco | 11360-070 |
| Pen Strep | Gibco | 15140-122 |
| L-Glutamine 200 mM | Gibco | A29168-01 |
| Vitamin | Gibco | 11120052 |
| PBS, 1X | Corning | 21-040-CV |
| 0.25% Trypsin-EDTA | Gibco | 25200-056 |
| Accutase | Corning | 25-058-CI |
| 70% V/V Ethanol | Fisher Bioreagents | BP8201-4 |
| Absolute Ethanol (200 proof) | Fisher Bioreagents | BP2818-4 |
| **RT-PCR Reagents** | **Manufacturer** | **Catalog #** |
| TRIzol reagent | Ambion | 15596018 |
| Chloroform | Fisher Scientific | C298-500 |
| Isopropyl Alcohol | Sigma | 19516-500ML |
| RNeasy Mini Kit (RNA isolation kit) | Qiagen | 74104 |
| Ambion DEPC-Treated Water | Invitrogen | AM9915G |
| iScript cDNA Synthesis Kit | Bio-Rad | 1708891 |
| SYBR Green Master mix | Applied Biosystems | 4309155 |
| TE Buffer pH 8.0 | Integrated DNA Technologies | 11-05-01-13 |
| **PAGE-Western Reagents** | **Manufacturer** | **Catalog #** |
| Cell lysis Buffer (10X) | Cell Signaling Technology | 9803 |
| RIPA Buffer (10X) | Cell Signaling Technology | 9806 |
| cOmplete Mini (protease inhibitor cocktail tablets) | Roche Diagnostics GmbH | 11836153001 |
| Phosphatase Inhibitor Cocktail-2 | Sigma-Aldrich | P5726-5ML |
| Phosphatase Inhibitor Cocktail-3 | Sigma-Aldrich | P0044-5ML |
| Phenylmethylsulfonyl fluoride (PMSF) | Cell Signaling Technology | 8553 |
| Precision Plus Protein Kaleidiscope | Bio-Rad | 1610375 |
| Laemmli SDS sample buffer, reducing (4X) | Thermo Scientific | J60015.AD |
| 10x Tris/Glycine/SDS Buffer | Bio-Rad | 1610772 |
| Albumin, Bovine, Fraction V, 35% soln. | Thermo Scientific | J64248.AE |
| Tris-buffered saline (TBS, 20X) pH 7.4 | Thermo Scientific | J60877.K3 |
| Tween 20 | Thermo Scientific | J20605-AP |
| Mini-Protean TGX Gels | Bio-Rad | 4561086 |
| Precision Plus Protein Standards | Bio-Rad | 1610375 |
| SuperSignal West Pico PLUS Chemiluminescent Substrate | Thermo Scientific | 34578 |
| SuperSignal West Femto Maximum Sensitivity Substrate | Thermo Scientific | 34096 |
| **T cell Isolation/ Exhaustion/ Cytotoxicity Reagents** | **Manufacturer** | **Catalog #** |
| Hoechst 33342 | Invitrogen | H1399 |
| Tumor dissociation Kit, Mouse | Miltenyi Biotec |  |
| Red Blood Cell Lysis Solution, 10X | Miltenyi Biotec | 130-094-183 |
| CD8 TIL Microbeads, Mouse (CD8+ T cell isolation Kit) | Miltenyi Biotec | 130-116-478 |
| MACS -BSA Stock Solution | Miltenyi Biotec | 130-091-376 |
| autoMACS Rinsing Solution | Miltenyi Biotec | 130-091-222 |
| QuadroMACS magnetic bead separation system | Miltenyi Biotec | 130-091-051 |
| LS Column | Miltenyi Biotec | 130-042-401 |
| Mouse IL-2 | Elabscience | PKSM041320 |
| Anti-CD3ε antibody | Bio cell | BE0001-1 |
| Anti-CD28 antibody | Bio cell | BE0015-5 |
| **Western Blot Antibodies** | **Manufacturer** | **Catalog #** |
| HIF-1α antibody | Cell Signaling Technology | 14179 |
| VEGFA antibody | Abcam | AB46154 |
| PD-L1 antibody | R&D Systems | MAB90781 |
| C-Myc antibody | Cell Signaling Technology | 13987 |
| pSTAT3 antibody | Cell Signaling Technology | 9134 |
| α-Tubulin antibody | Cell Signaling Technology | 3873 |
| Anti-mouse IgG, HRP-linked | Cell Signaling Technology | 7076 |
| Anti-mouse IgG, HRP conjugated | R&D Systems | HAF008 |
| **Flow Reagents** | **Manufacturer** | **Catalog #** |
| Zombie Violet™ Fixable Viability Kit | BioLegend | 423113 |
| True-Stain Multi-Fluor Buffer | BioLegend | 426105 |
| True-Stain Monocyte Blocker | BioLegend | 426102 |
| Fc Block | BioLegend | 156603 |
| eBioscience™ Foxp3 / Transcription Factor Staining Buffer Set | Invitrogen | 00-5523-00 |
| UltraComp eBeads™ Plus Compensation Beads | Invitrogen | 01-3333-41 |
| ArC™ Amine Reactive Compensation Bead Kit | Invitrogen | A10346 |
| **Immunohistochemistry Reagents** | **Manufacturer** | **Catalog #** |
| Xylenes | Fisher Chemical^TM^ | X5-4 |
| Antigen Unmasking Solution, Tris-Based | Vector Laboratories | H-3301-250 |
| 30% Hydrogen Peroxide | Fisher Chemical^TM^ | H325-500 |
| Normal Goat Serum | Cell Signaling Technology | 5425S |
| CD31 (PECAM-1) Rabbit mAb | Cell Signaling Technology | 77699S |
| SignalStain Boost IHC Detection Reagent (HRP, Rabbit) | Cell Signaling Technology | 8114S |
| DAB Substrate | Vector Laboratories | SK-4105 |
| Hematoxylin | Cell Signaling Technology | 14166S |
| Acetic Acid, Glacial | Fisher Chemical^TM^ | A38-500 |
| Ammonium hydroxide (NH_4_OH) 30% | Sigma-Aldrich | 05002-1L |
| Cytoseal^TM^ 60 Mountant | Epredia | 8310-4 |
