## Supplemental Table 2 for "mTOR inhibition augments antitumor immune effector response by reprogramming the *TP53*-mutant, immune-cold HNSCC tumor microenvironment"

**Supplementary Table S-2.**

| **Panel** | **Marker/Fluorophore** | **Manufacturer** | **Catalog #** | **Dilution used** |
| --- | --- | --- | --- | --- |
| Lymphoid Panel | Zombie Violet | BioLegend | 423113 | 500 |
|  | CD45 AF700 | BioLegend | 103128 | 2000 |
|  | CD19 BV711 | BioLegend | 115555 | 400 |
|  | NK1.1 BV605 | BioLegend | 108753 | 100 |
|  | CD3 APC/Fire 750 | BioLegend | 100248 | 100 |
|  | CD4 BV785 | BioLegend | 100453 | 400 |
|  | CD8 BV650 | BioLegend | 100742 | 400 |
|  | CD25 PE | BioLegend | 102008 | 200 |
|  | CD62L PE/Cy7 | BioLegend | 104418 | 800 |
|  | PD1 PE-Dazzle 594 | BioLegend | 109116 | 200 |
|  | FoxP3 APC | eBioscience | 17-5773-82 | 100 |
| Myeloid Panel | Zombie Violet | BioLegend | 423113 | 500 |
|  | CD45 AF700 | BioLegend | 103128 | 2000 |
|  | CD11b AF488 | BioLegend | 101217 | 2000 |
|  | CD11c APC | BioLegend | 117310 | 400 |
|  | F4/80 BV650 | BioLegend | 123149 | 100 |
|  | GR-1 BV711 | BioLegend | 108443 | 200 |
|  | Ly6C PE/Cy7 | BioLegend | 128017 | 800 |
|  | I-A/I-E (MHCII) PE | Biolegend | 107607 | 800 |
| CD8+ IFN-γ+ Detection Panel | Zombie Violet | BioLegend | 423113 | 500 |
|  | CD45 AF700 | BioLegend | 103128 | 2000 |
|  | CD19 BV711 | BioLegend | 115555 | 400 |
|  | CD3 APC/Fire 750 | BioLegend | 100248 | 100 |
|  | CD4 BV785 | BioLegend | 100453 | 400 |
|  | CD8 BV650 | BioLegend | 100742 | 400 |
|  | IFN-γ BV605 | Biolegend | 505840 | 100 |
| PD-1 expressing T cell Detection Panel | Zombie Violet | BioLegend | 423113 | 500 |
|  | CD45 AF700 | BioLegend | 103128 | 2000 |
|  | CD3 APC/Fire 750 | BioLegend | 100248 | 100 |
|  | CD4 BV785 | BioLegend | 100453 | 400 |
|  | CD8 BV650 | BioLegend | 100742 | 400 |
|  | PD1 PE-Dazzle 594 | BioLegend | 109116 | 200 |
