## Supplementary figures and images for "mTOR inhibition augments antitumor immune effector response by reprogramming the *TP53*-mutant, immune-cold HNSCC tumor microenvironment"

### Supplemental Figure 3

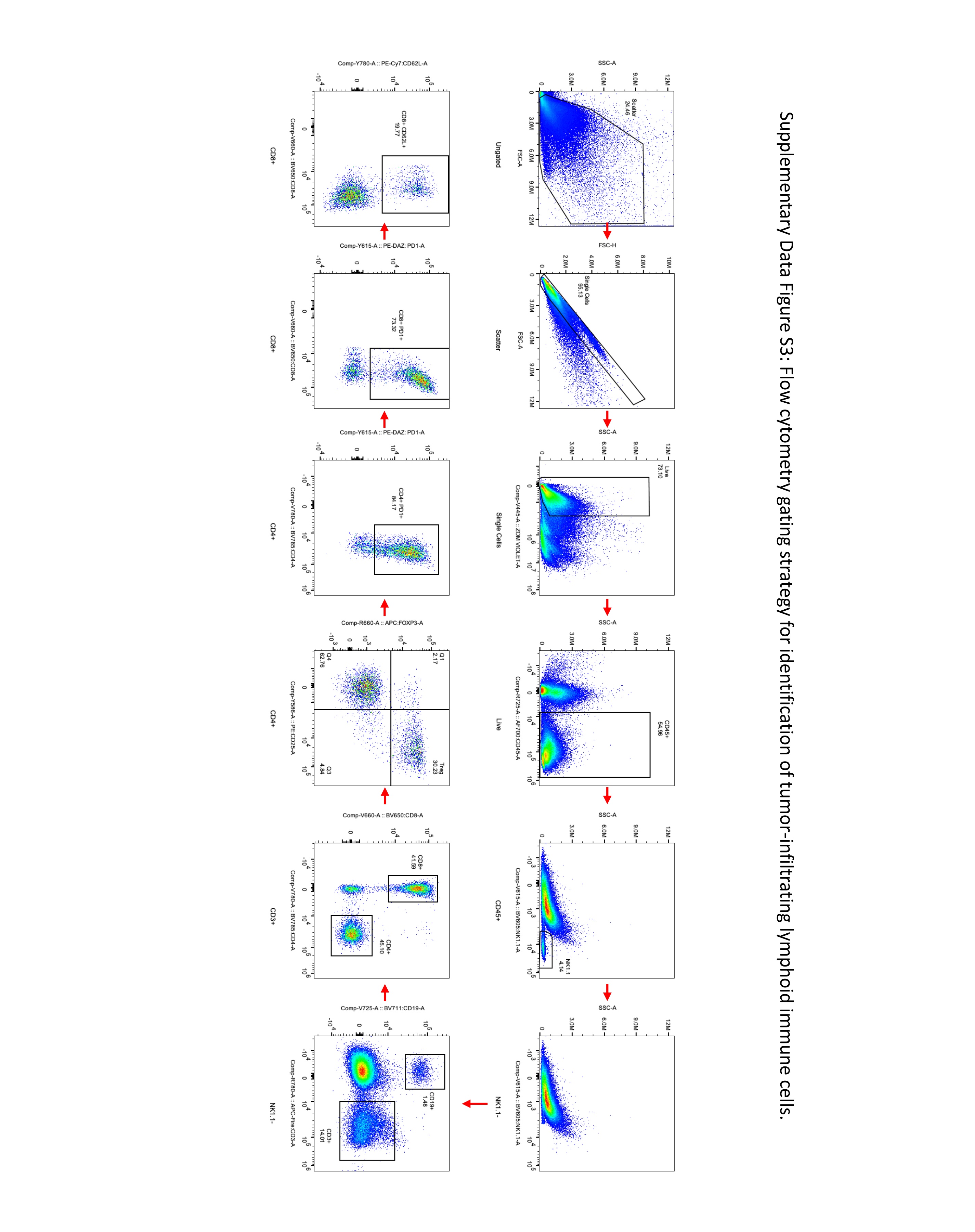

### Supplemental Figure 4

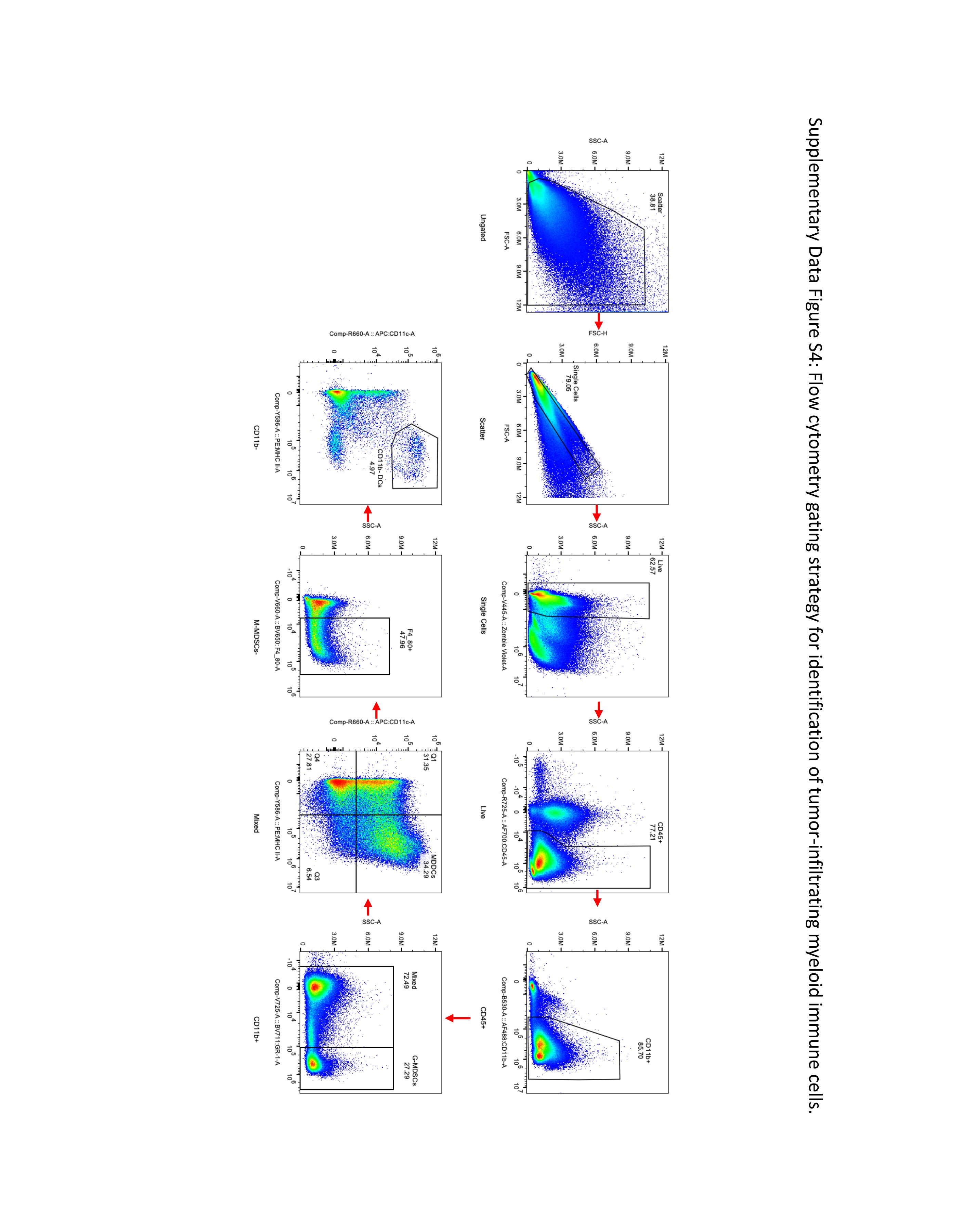

### Supplemental Figure 5

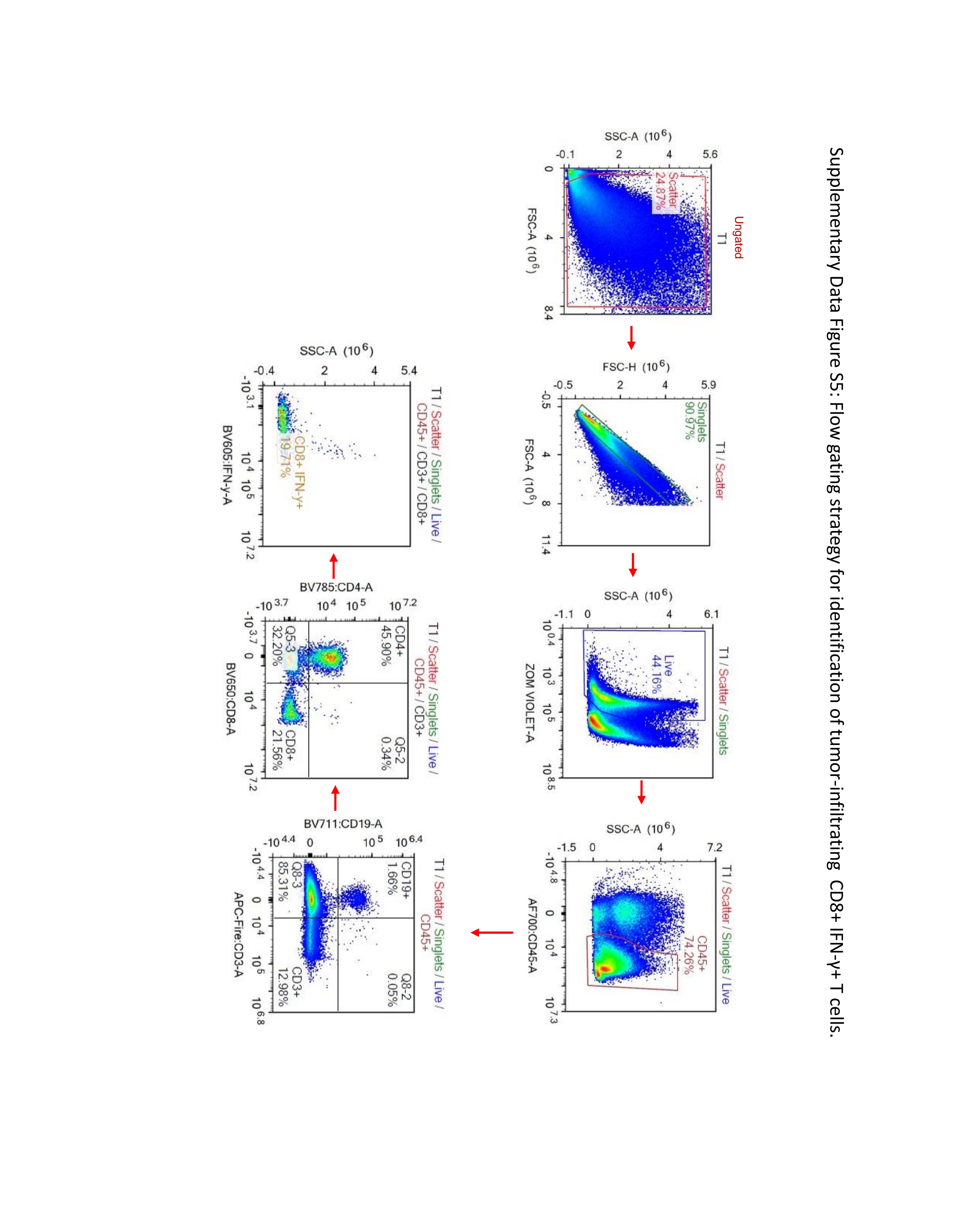

### Supplemental Figure 7

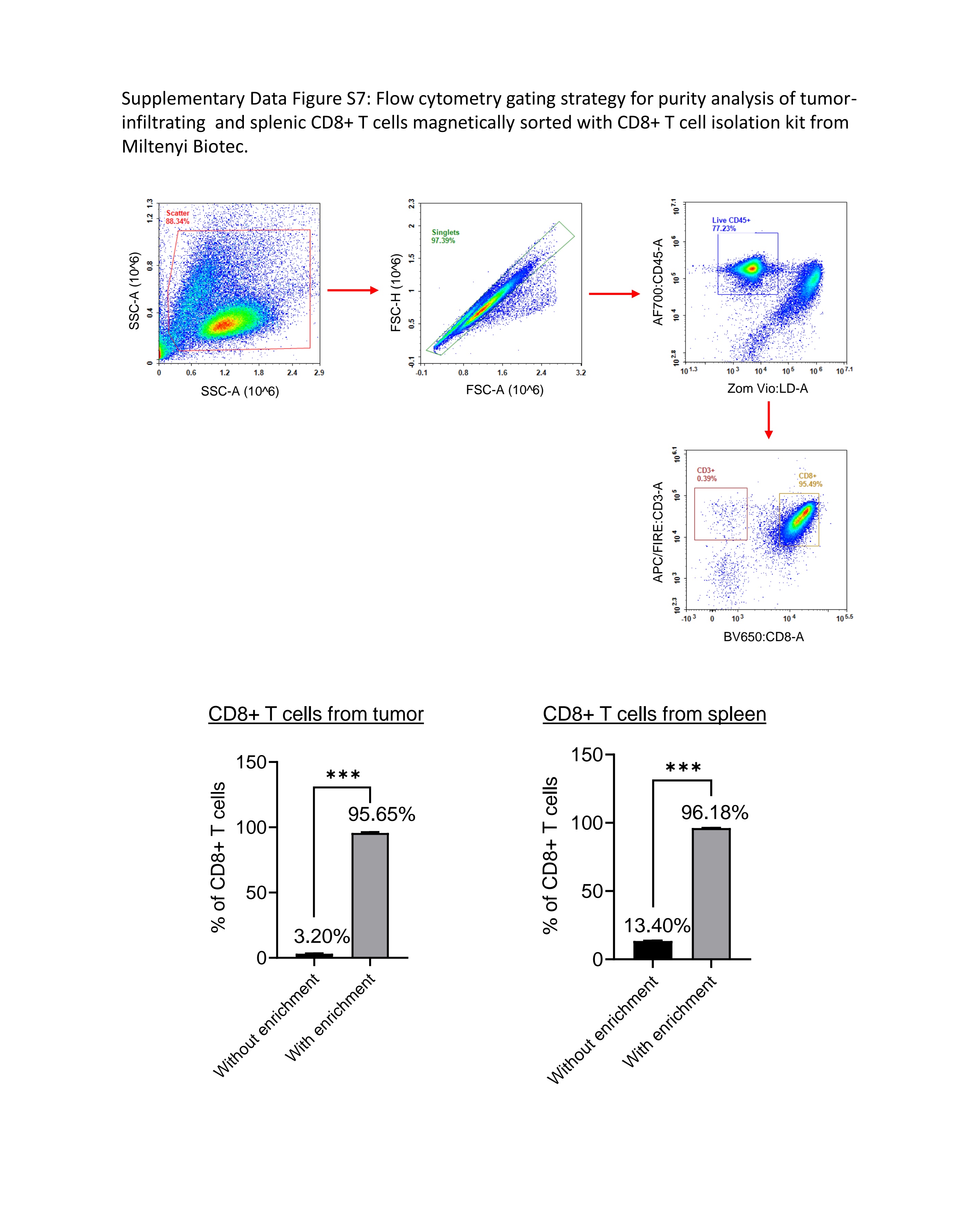

### Supplemental Figure 8

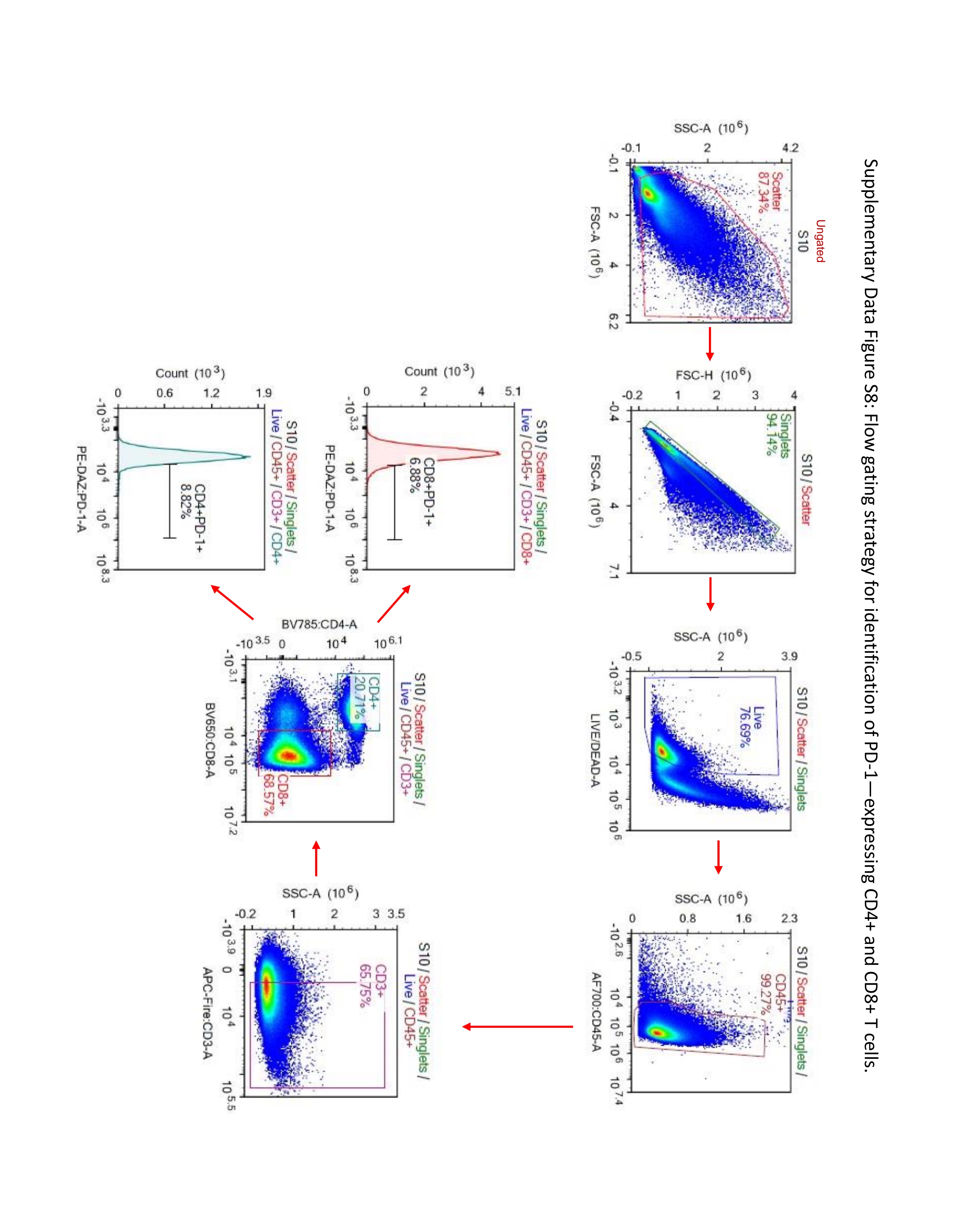
