## Supplemental Table 6 for "mTOR inhibition augments antitumor immune effector response by reprogramming the *TP53*-mutant, immune-cold HNSCC tumor microenvironment"

**Supplementary Table S6 (PCR Primers)**

| **Gene Name** | **Forward primer sequence**  **(5'->3')** | **Reverse primer sequence**  **(5'->3')** | **Gene Bank Accession** |
| --- | --- | --- | --- |
| CXCL10 | TGAGAGACATCCCGAGCCAA | GAGGCAGAAAATGACGGCAG | NM_021274 |
| TNF-α | CAGGCGGTGCCTATGTCTC | CGATCACCCCGAAGTTCAGTAG | NM_013693 |
| IRF1 | ATGCCAATCACTCGAATGCG | TTGTATCGGCCTGTGTGAATG | NM_001159396 |
| GZMB (Granzyme B) | TCTCGACCCTACATGGCCTTA | TCCTGTTCTTTGATGTTGTGGG | NM_013542 |
